# Hematopoietic-SLC37A2 deficiency accelerates atherosclerosis in LDL receptor-deficient mice

**DOI:** 10.1101/2021.08.06.455449

**Authors:** Qingxia Zhao, Zhan Wang, Allison K. Meyers, Jennifer Madenspacher, Manal Zabalawi, Elena Boudyguina, Fang-Chi Hsu, Charles M. McCall, Cristina M. Furdui, John S. Parks, Michael B. Fessler, Xuewei Zhu

## Abstract

Macrophages play a central role in the pathogenesis of atherosclerosis. Our previous study demonstrated that solute carrier family 37 member 2 (SLC37A2), an endoplasmic reticulum-anchored phosphate-linked glucose-6-phosphate transporter, negatively regulates macrophage Toll-like receptor activation by fine-tuning glycolytic reprogramming *in vitro*. Whether macrophage SLC37A2 impacts *in vivo* macrophage inflammation and atherosclerosis under hyperlipidemic conditions is unknown. We generated hematopoietic cell-specific SLC37A2 knockout and control mice in C57Bl/6 Ldlr^-/-^ mice by bone marrow transplantation. Hematopoietic-specific SLC37A2 deletion in Ldlr^-/-^ mice increased plasma lipid concentrations 12-16 wks of Western diet induction, attenuated macrophage anti-inflammatory responses, and resulted in more atherosclerosis compared to Ldlr^-/-^ mice transplanted with wild type bone marrow. Aortic root intimal area was inversely correlated with plasma IL-10 levels, but not total cholesterol concentrations, suggesting inflammation but not plasma cholesterol was responsible for increased atherosclerosis in bone marrow SLC37A2-deficient mice. Our *in vitro* study demonstrated that SLC37A2 deficiency impaired IL-4-induced macrophage activation, independently of glycolysis or mitochondrial respiration. Importantly, SLC37A2 deficiency impaired apoptotic cell-induced glycolysis, subsequently attenuating IL-10 production. Our study suggests that SLC37A2 expression is required to support alternative macrophage activation *in vitro* and *in vivo. In vivo* disruption of hematopoietic SLC37A2 accelerates atherosclerosis under hyperlipidemic pro-atherogenic conditions.

## Introduction

Atherosclerosis is driven by hyperlipidemia and exacerbated by chronic low-grade inflammation ^1-6^. Macrophages are among the most abundant immune cells within atherosclerotic plaques ^1,5,7-10^. Inside the atherosclerotic plaque, macrophages take up modified LDL (i.e., oxidized LDL; oxLDL), forming lipid laden-macrophages or foam cells, which worsen inflammation and promote atherosclerotic plaque growth ^1,5,7-10^. As a heterogeneous cell population, macrophages can be roughly grouped into two categories based on their inflammatory activities: classically activated pro-inflammatory (M1) macrophages and alternatively activated anti-inflammatory macrophages ^11^. The anti-inflammatory (M2) macrophages can be subdivided into M2a, 2b, 2c, and 2d based on the stimuli and resultant transcriptional changes ^11^. These *in vitro* models of macrophage polarization (M1 vs. M2) are more simplified than the microenvironment that macrophages encounter in an atherosclerotic lesion ^12^. Nevertheless, mounting evidence suggests that macrophage plasticity or phenotypic switch impacts atherosclerotic lesion progression and regression ^1,13,14^. Uncovering the underlying mechanisms governing activation and deactivation of macrophages and their phenotypic switch provides a promising avenue to prevent and treat atherosclerosis.

Macrophages rewire intracellular metabolic pathways upon activation, which in turn modify their cellular function ^15-17^. Pro-inflammatory macrophages, such as lipopolysaccharide (LPS)-stimulated macrophages (M(LPS)), requires glycolysis to mount an effective inflammatory response. However, whether glycolytic reprogramming is necessary for alternative macrophage activation is still debatable and requires further investigation. Moreover, little is known about how and to what extent macrophages reprogram cellular metabolism in response to microenvironmental stimuli to promote or resolve inflammation under pro-atherogenic conditions. A positive association between glycolysis and plaque macrophage inflammation was documented as increased glucose metabolic activity in human symptomatic and unstable plaque macrophages compared with asymptomatic lesions ^18^. However, macrophage glycolytic rate does not always influence atherogenesis, at least based on animal studies. For example, myeloid deletion of glucose transporter 1 (GLUT1), the primary glucose transporter on the plasma membrane in macrophages, does not alter atherosclerotic plaque size or macrophage content in Ldlr^-/-^ mice ^19^. Moreover, myeloid overexpression of GLUT1 increases glucose flux in macrophages and enhances macrophage inflammation *in vitro* but is insufficient to promote atherosclerosis ^20^. These findings suggest a context-dependent regulation of macrophage inflammation and disease development by cellular metabolic processes in acute vs. chronic low-grade inflammation.

Solute carrier family 37 member 2 (SLC37A2), an endoplasmic reticulum-anchored phosphate-linked glucose-6-phosphate transporter, is highly expressed in macrophages ^21-23^. We have recently reported that SLC37A2 plays a pivotal role in murine macrophage inflammatory activation and cellular metabolic rewiring ^24^. SLC37A2 deletion reprograms macrophages to a hyper-glycolytic state of energy metabolism and accelerates M (LPS) activation, partially depending on nicotinamide adenine dinucleotide (NAD^+^) biosynthesis. Blockade of glycolysis or the NAD^+^ salvage pathway normalizes the differential expression of pro-inflammatory cytokines between control and SLC37A2-deficient macrophages. Conversely, overexpression of SLC37A2 lowers macrophage glycolysis and significantly reduces LPS-induced pro-inflammatory cytokine expression. Our published work suggests that SLC37A2 is a negative regulator of murine macrophage pro-inflammatory activation by down-regulating glycolytic reprogramming.

Despite these findings, it remains unclear whether macrophage SLC37A2 impacts macrophage pro- or anti-inflammatory activation in vivo under pathologic conditions. Further, it is not known whether macrophage SLC37A2-mediated inflammation affects the pathogenesis of inflammatory diseases. Here, we found that SLC37A2 expression is necessary to maintain alternative macrophage activation *in vitro*. Our data suggest that SLC37A2 positively regulates IL-4-induced macrophage alternative activation, independent of glycolysis or mitochondrial respiration. SLC37A2 positively regulates apoptotic cell-induced macrophage alternative activation through modulation of glycolysis. Lastly, we found that disruption of hematopoietic SLC37A2 impairs anti-inflammatory responses and worsens hyperlipidemia-induced atherosclerosis in Ldlr^-/-^ mice.

## Materials and Methods

### Animals, bone marrow transplantation (BMT), and diet feeding

#### Animals

*Slc37a2* global knockout mice in the C57BL/6J background (T1837) were purchased from Deltagen, Inc ^24^. Heterozygous *Slc37a2* knockout mice were intercrossed to obtain wild type (WT) and homozygous knockout (*Slc37a2*^*-/-*^) mice. Ldlr^-/-^ (stock 002207) mice were purchased from Jackson Laboratories. Mice were housed in a pathogen-free facility on a 12 h light/dark cycle and received a standard laboratory diet.

#### BMT

7 × 10^6^ BM cells from female donor (WT and *Slc37a2*^*-/-*^) mice were injected into the retro-orbital venous plexus of irradiated male Ldlr^-/-^ recipient mice. Repopulation of blood leukocytes after BMT was evaluated after 16 wks of diet feeding by determining the percentage expression of the Y-chromosome-associated sex-determining region Y gene (Sry) in genomic DNA obtained from white blood cells, as described previously ^25^.

#### High-fat western diet (WD) feeding

After 5 wks for recovery from radiation, mice were switched from a standard laboratory diet to a high-fat WD containing 42% calories from fat and 0.2% cholesterol (TD. 88137, Teklad) for an additional 16 wks to induce advanced atherosclerosis.

All animal experimental protocols were approved by the Wake Forest University Animal Care and Use Committee.

### Analysis of atherosclerotic lesions

Aortic root atherosclerosis was assessed as described before ^25^. Briefly, aortic root sections were stained in 0.5% Oil Red O and counterstained with hematoxylin. To measure necrosis, boundary lines were drawn using NIH ImageJ software around regions that were free of H&E staining. We used a 3000 µm^2^ threshold to avoid counting regions that may not represent substantial areas of necrosis, as described before ^25^. Stained sections were photographed with an Olympus DP71 digital camera and quantified using NIH ImageJ software. Results were expressed as cross-sectional plaque area (H&E) or plaque necrosis (necrotic core), or percent of total plaque area or necrosis core. Macrophages and T cells in aortic root were stained by incubating aortic root cross-sections with the primary antibodies to CD68 (Bio-Rad) and CD3 (Abcam), followed by the biotinylated secondary antibody. The staining was visualized using the ABC reagent (ABC vector kit; Vector) and DAB substrate chromogen (Dako). Apoptotic cells in atherosclerotic lesions were stained using the Click-iT Plus TUNEL Assay kit (Thermo Fisher) according to the manufacturer’s protocol. The sections were then stained with CD68 antibody (Bio-Rad), and nuclei were stained with DAPI. Only TUNEL-positive cells that co-localized with DAPI-positive nuclei were counted as apoptotic cells. Efferocytosis was determined by counting the number of macrophage-associated versus free apoptotic cells, following established methods published before ^25-29^. Aortic root cross-sections were also stained with Masson’s Trichrome stain to quantify collagen deposition. Areas stained blue within lesions were identified as collagen-positive using NIH ImageJ software.

### Cell culture and treatment

#### Peritoneal macrophage culture

Peritoneal cells were harvested from mice after 16-wk diet feeding ^30,31^. Cells were plated in RPMI-1640 media containing 100 U/ml penicillin and 100 μg/ml streptomycin. After a 2 h incubation, floating cells were removed by washing with PBS, and adherent macrophages were lysed using Trizol (Invitrogen) for RNA extraction.

#### BMDM culture

Mouse bone marrow was cultured in low glucose DMEM supplemented with 30% L929 cell-conditioned medium and 20% FBS for 6-7 days until the cells reached confluence. BMDMs were then reseeded in culture dishes overnight in RPMI 1640 medium containing 1% Nutridoma-SP medium (Sigma-Aldrich) before any treatment ^24^.

#### Jurkat T cell culture

Jurkat T cells (TIB-152, ATCC) were maintained in 10% FBS containing RPMI-1640 medium. To induce apoptosis, Jurkat T cells (2×10^6^/ml) were treated with 1 μM staurosporine in RPMI-1640 media containing 100 U/ml penicillin and 100 μg/ml streptomycin for 4 h.

#### Macrophage stimulation

BMDMs were incubated with 20 ng/ml IL-4 or ACs (ratio of ACs to macrophages was 5:1) in 10% FBS containing for indicated times as written in the figure legends. To induce foam cell formation, BMDMs were treated with 25 or 50 μg/ml oxLDL (Athens Research & Technology) for 0-24 h. In some experiments, BMDMs were pretreated for 30 min with hexokinase inhibitor 2-deoxy-D-glucose (2-DG; 10 mM, Sigma-Aldrich), actin polymerization inhibitor cytochalasin D (10 μM, Cayman), or fatty acid β-oxidation inhibitor etomoxir (50 μM, Cayman) and subsequently treated with apoptotic cells for an additional 4 h in the presence of each inhibitor.

### Cytokine quantification

Total RNA in tissues or macrophages was extracted using Trizol (Invitrogen). cDNA preparation and real-time PCR were conducted as described previously ^24,25^. Primers were listed in Table S1. Concentrations of cytokines/chemokines in plasma, liver homogenate, or cell culture supernatant were measured using Bioplex assay or ELISA according to the manufacturer’s instructions.

### Flow cytometry

Peripheral blood and splenocytes were stained with Gr1(Ly6C/G)-PerCP-Cy5.5 (BD Pharmingen, cat# 552093), CD115-APC (eBioscience, cat# 17-1152-82), CD11b-APC Cy7 (BD Bioscience, cat# 557657), and V450-CD45 (BD Horizon, cat# 560501) ^30^. Data were acquired on a BD FACS Canto II instrument (BD Biosciences) and analyzed using FACSDiva software v6.1.3 (BD Biosciences). To examine the apoptotic rate, Jurkat T cells were stained with Annexin V-APC and PI (Thermo Fisher). Data were acquired on a BD FACSCalibur (BD Biosciences) and analyzed using Flowjo V7.6.5 (BD Biosciences).

### Glucose tolerance tests and insulin tolerance tests

Mice were fasted overnight before intraperitoneal injection of 1 g glucose/kg BW. One wk later, the same groups of mice were used for intraperitoneal injection of 0.75 U of regular human insulin/kg BW after a 4 h fast. Blood glucose concentrations were measured at 0, 15, 30, 60, and 120 min after each injection ^31^.

### Lipid analysis

4 h-fasting plasma or liver total cholesterol (TC), free cholesterol (FC) (Wako), and triglyceride (TG) (Roche) were determined by enzymatic analysis. Plasma was fractionated by FPLC to determine cholesterol distribution among the lipoprotein classes ^32^. Liver lipids were normalized to wet liver weight ^25^. Macrophage cholesterol content was measured by gas-liquid chromatography ^33^ and normalized to cellular protein ^33^.

### Seahorse assay

2 × 10^5^ BMDMs were plated into each well of Seahorse XF96 cell culture microplates (Agilent Technologies) and cultured overnight before treated with or without 20 ng/ml IL-4 for 6 or 24 h or ACs for 3 h. Basal and IL-4- or AC-induced changes in oxygen consumption rate (OCR) and extracellular acidification rate (ECAR) in BMDMs were measured with a Seahorse XF96 Extracellular Flux Analyzer (Agilent Technologies) under basal conditions and following the sequential addition of 10 mM glucose, 1 µM oligomycin, 1.5 µM fluoro-carbonyl cyanide phenylhydrazone (FCCP), 100 nM rotenone plus 1 µM antimycin A, or 50 mM 2DG (all the compounds were from Agilent Technologies), as described in the figure legends. After the assay, 3 μM Hoechst (Life Technologies) was added to each well to stain nuclei for cell counting. Results were collected with Wave software version 2.6 (Agilent Technologies). Data was normalized to cell numbers.

### Western blotting

BMDM protein concentration was measured using the BCA protein assay kit (Pierce). Rabbit anti-SLC37A2 polyclonal antibody was made against the peptide CTPPRHHDDPEKEQDNPEDPVNSPYSSRES (LAMPIRE Biological Lab Inc.) and used at a dilution of 1:500 ^24^.

### Statistics

Statistical analysis was performed using GraphPad Prism software 7 (GraphPad Software) except for the plasma lipid analysis in Figures 1A-1D. Data are presented as the mean ± SEM unless indicated otherwise. Differences were compared with Student’s t-test or two-way ANOVA with post hoc Tukey’s multiple comparison test as indicated in the figure legends. In Figures 1A-1D, the mixed-effects models were used to compare lipids variables between genotype groups at each time point. The use of random intercepts provided a source of autocorrelation between repeated measures. Genotype groups and weeks and the interaction between genotype groups and weeks were included in the model. Contrasts were used to compare lipid variables at each measured week. P < 0.05 was considered statistically significant.

**Figure 1.**
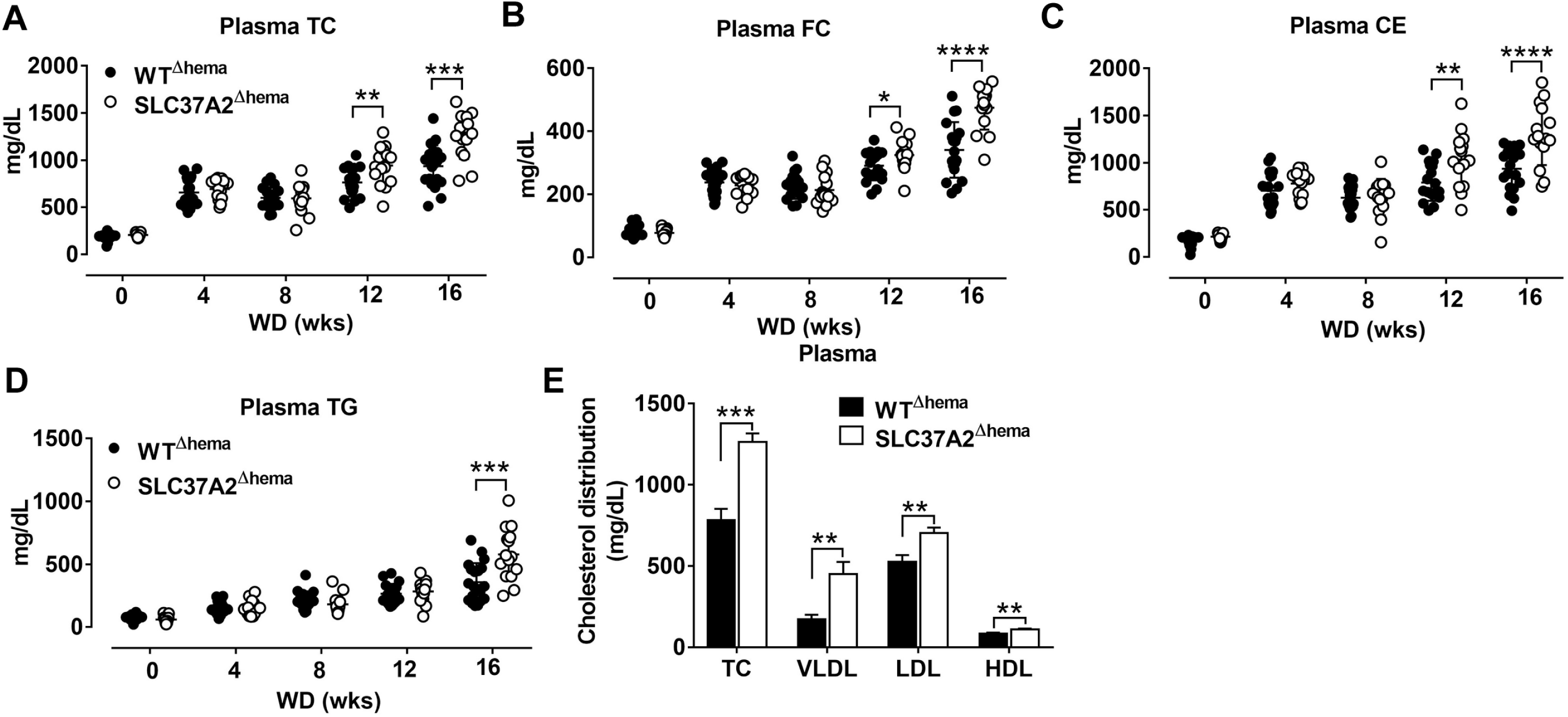
Hematopoietic SLC37A2 deletion increases plasma lipids in WD-fed Ldlr^-/-^ mice. Hematopoietic SLC37A2 knockout (SLC37A2^Δhema^) and control (WT^Δhema^) Ldlr^-/-^ mice were switched from a standard laboratory diet to high-fat WD for 16 wks to promote the development of advanced atherosclerosis. (**A-D**) Fasting (4 h) plasma total cholesterol (TC), free cholesterol (FC), cholesterol ester (CE), and triglyceride (TG) concentrations were measured by enzymatic assays over time (0–16 wks) (n=12-16). (**E**) Plasma cholesterol distribution among lipoproteins after 16 wks of WD feeding was determined after fractionation of plasma by fast protein liquid chromatography (n=6-8). Data are expressed as mean ± SEM. ^**^ P<0.01, ^***^ P<0.001, ^****^ P<0.0001, unpaired, two-tailed Student’s t-test. HDL indicates high-density lipoprotein; LDL, low-density lipoprotein; and VLDL, very-LDL.

## Results

### Hematopoietic SLC37A2 deletion increases plasma lipids in WD-fed Ldlr^-/-^ mice

We previously showed that SLC372 is a novel regulator of macrophage inflammation by controlling glycolysis ^24^. In this study, we wanted to know whether mice lacking SLC37A2 in macrophages were at an increased risk of developing atherosclerosis. To do this, we generated hematopoietic SLC37A2 knockout (SLC37A2^Δhema^) mice in Ldlr^-/-^ background by BMT. The BMT efficiency was ∼90% based on the male Sry gene expression in blood leukocytes isolated from recipient mice examined after 16-wk diet feeding (**Figures S1A** and **S1B**). Four wks of WD feeding increased plasma lipid (including TPC, FC, CE, and TG) concentrations in both genotypes. Interestingly, SLC37A2^Δhema^ mice displayed significantly higher plasma cholesterol concentrations after 12 and 16 wks of diet feeding (**Figures 1A-1C**). SLC37A2^Δhema^ mice also showed significantly higher plasma TG concentration after 16 wks of diet feeding (**Figure 1D**). Furthermore, we observed significantly higher plasma VLDL and LDL cholesterol concentrations but only a marginal increase in HDL cholesterol in SLC37A2^Δhema^ vs. WT mice after 16 wks of diet feeding (**Figure 1E**). Collectively, these results suggest a novel role for hematopoietic SLC37A2 in lipid metabolism under pro-atherogenic conditions.

### Hematopoietic SLC37A2 deletion increases liver CE and reduces hepatic macrophage activation in WD-fed Ldlr^-/-^ mice

Given the increased plasma lipid concentrations in high-fat WD-fed SLC37A2^Δhema^ mice, we next measured liver lipid concentrations. Consistent with increased plasma lipid concentrations, we observed a significant increase in hepatic TC and CE, but not FC or TG in SLC37A2^Δhema^ vs. WT mice (**Figure 2A**). The increased plasma and liver lipid concentrations in SLC37A2^Δhema^ mice prompted us to examine the hepatic expression of genes involved in lipid metabolism in diet-fed mice. As expected, Slc37a2 transcript was significantly lower in SLC37A2^Δhema^ mice receiving bone marrow from *Slc37a2*^*-/-*^ mice (**Figure 2B**). We found that hepatic expression of genes responsible for de novo lipogenesis was similar between genotypes of mice (**Figure 2C**). There was a slightly but significantly decreased expression of cholesterol transporter protein ABCA1 (**Figure 2C**). Interestingly, hepatic expression of the receptor for fatty acid uptake (CD36), enzymes for fatty acid oxidation (Cpt1a, Acox1, Acadl), and transcriptional factors and coactivators regulating fatty acid oxidation (PGC1α, PGC1β, PPARδ, and PPARγ) were all down-regulated in SLC37A2^Δhema^ vs. WT liver (**Figure 2D**), suggesting an impaired fatty acid utilization in SLC37A2^Δhema^ mouse liver at least at the transcriptional level. Because alternative activation of hepatic macrophages promotes liver fatty acid oxidation and improves metabolic syndrome ^34^, we next examined macrophage pro- and anti-inflammatory states by measuring hepatic gene expression of macrophage markers, M1-type cytokines, and chemokines, and M2 macrophage markers. Different from our *in vitro* study, in which we observed increased pro-inflammatory cytokine production in SLC37A2^Δhema^ macrophages in response to TLR activation ^24^, under pro-atherogenic conditions, hematopoietic SLC37A2 deficiency has a minor effect on the expression of genes encoding macrophage markers (**Figure 2E**) and pro-inflammatory cytokines and chemokines (**Figures 2F** and **2G**). Rather, we observed a slightly decreased IFN-γ and MCP-1 (**Figures 2F** and **2G**) protein concentration in SLC37A2^Δhema^ liver, suggesting that SLC37A2 deletion in bone marrow is not sufficient to induce a pro-inflammatory response in the liver under pro-atherogenic conditions. Despite the equivalent or slightly decreased pro-inflammatory cytokines/chemokines in SLC37A2^Δhema^ liver, hepatic anti-inflammatory markers, including Arg1, Mrc1, Ym1, and IL-10, showed significantly lower expression in SLC37A2^Δhema^ vs. WT liver (**Figure 2H**). Consistent with the decreased transcript expression, hepatic IL-10 protein concentration showed a trend toward a decrease in SLC37A2^Δhema^ liver, relative to control (**Figure 2I**). Taken together, our results suggest that genetic deletion of SLC37A2 in bone marrow cells significantly impairs alternative activation of hepatic macrophages, associated with lower hepatic fatty acid oxidation gene expression and increased liver lipid accumulation.

**Figure 2.**
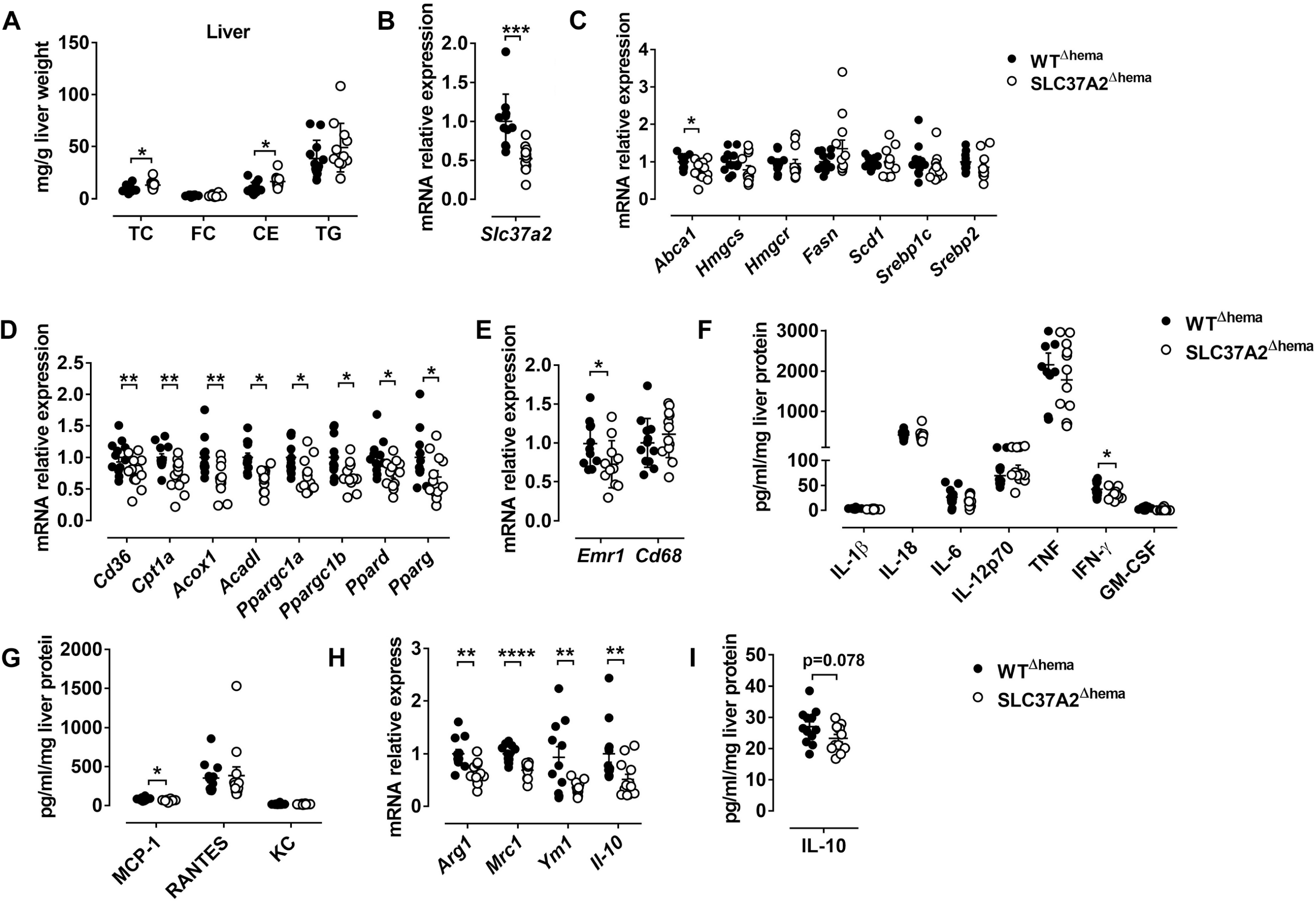
Hematopoietic SLC37A2 deletion increases liver CE and reduces hepatic macrophage activation in WD-fed LDLrKO mice. Irradiated Ldlr^−/−^ mice transplanted with WT or SLC37A2KO bone marrow were fed a high-fat western diet for 16 wks. (**A**) Fasting (4 h) total cholesterol (TC), free cholesterol (FC), cholesterol ester (CE), and triglyceride (TG) concentrations in the liver were measured by enzymatic assays (n=12 per genotype). (**B**) Slc37a2 transcript expression in liver (n=12 per genotype). (**C**) Relative transcript levels for cholesterol efflux, cholesterol synthesis, fatty acid synthesis, and transcriptional regulators controlling these pathways in the liver (n=12 per genotype). (**D**) Relative transcript levels for genes encoding key enzymes in fatty acid uptake, β-oxidation, and transcriptional regulators controlling these pathways in the liver (n=12 per genotype). (**E**) Relative transcript levels for genes encoding macrophage markers in the liver (n=12 per genotype). (**F-G**) Liver cytokine and chemokine protein concentrations (n=12 per genotype). (**H**) Relative transcript levels for genes encoding macrophage markers (E) and pro-inflammatory cytokines (F) in the liver (n=12 per genotype). (**I**) Liver IL-10 protein concentrations (n=12 per genotype). Data are expressed as mean ± SEM. ^***^ P<0.01, ^***^ P<0.001, ^****^ P<0.0001, unpaired, two-tailed Student’s t-test.

### Hematopoietic SLC37A2 deletion primarily impairs anti-inflammatory responses in WD-fed Ldlr^-/-^ mice

To determine whether hematopoietic SLC37A2 deletion affects inflammation at the systemic level, we first examined plasma concentrations of pro-inflammatory cytokines, chemokines, and anti-inflammatory cytokine using a multiplex assay. Consistent with the expression pattern of hepatic cytokines and chemokines, plasma pro-inflammatory cytokine concentrations did not differ between genotypes (**Figure 3A**). But, plasma MCP-1 concentration showed a 28.6% reduction in the SLC37A2^Δhema^ vs. WT mice (**Figure 3B**). Strikingly, the SLC37A2^Δhema^ mice displayed a 70% reduction of the anti-inflammatory cytokine IL-10 in plasma (**Figure 3C**) relative to their WT counterparts, suggesting hematopoietic SLC37A2 deletion impairs IL-10 production under pro-atherogenic conditions. Moreover, SLC37A2^Δhema^ vs. WT mice showed indistinguishable expression of pro-inflammatory cytokines (**Figure 3D**) but significantly attenuated Arg1 and Mrc1 (alternative activation markers) expression in resident peritoneal macrophages (**Figure 3E)** after 16-wk diet feeding. Together, our results suggest that hematopoietic SLC37A2 deletion primarily impairs anti-inflammatory responses in Ldlr^-/-^ mice when challenged with WD diet.

**Figure 3.**
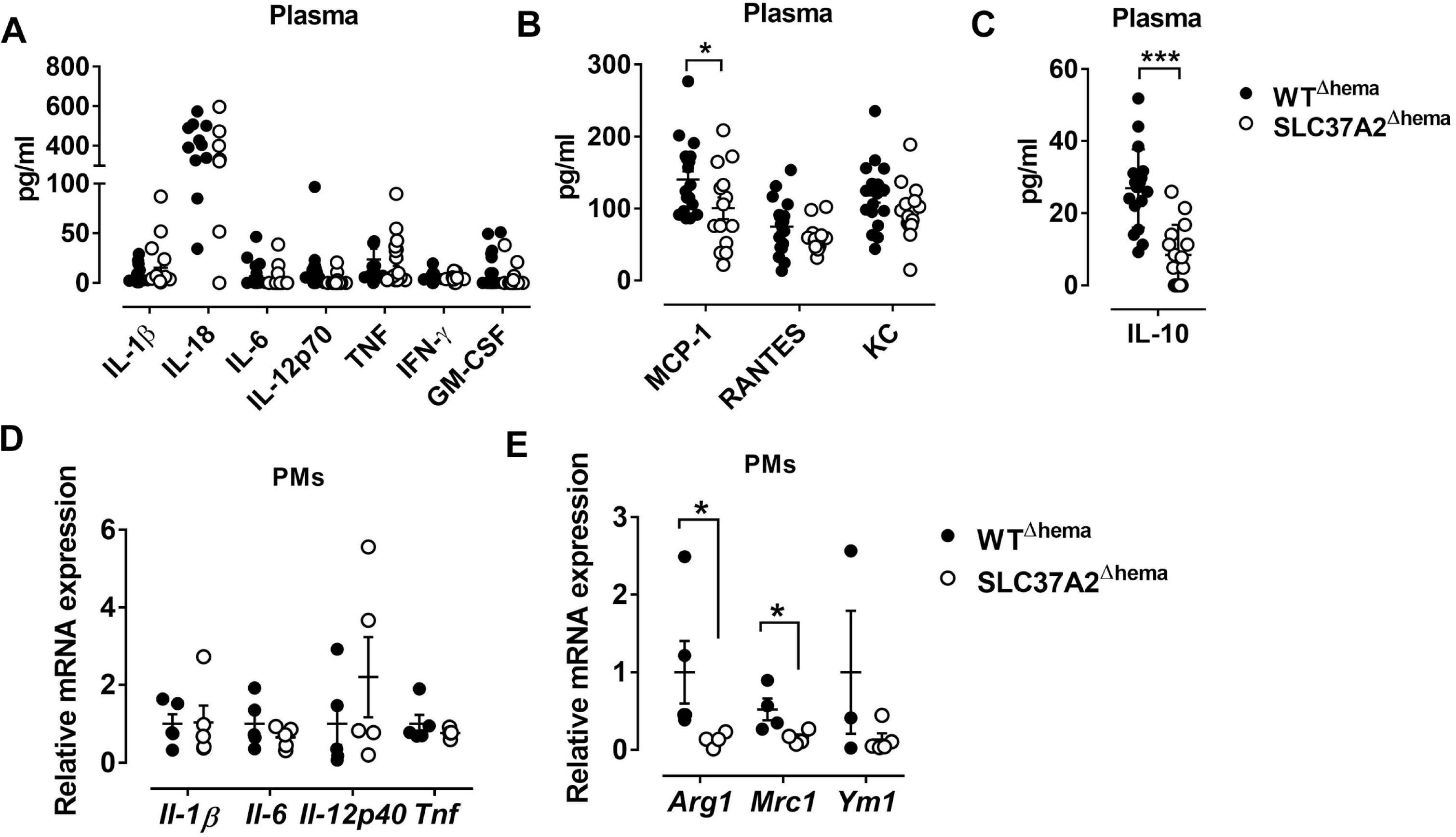
Hematopoietic SLC37A2 deletion primarily impairs anti-inflammatory responses in WD-fed Ldlr^-/-^ mice. Hematopoietic SLC37A2 knockout (SLC37A2^Δhema^) and control (WT^Δhema^) Ldlr^-/-^ mice were fed a high-fat western diet for 16 wks. (**A-C**) Plasma pro-inflammatory cytokine (A), chemokine (B), and anti-inflammatory cytokine IL-10 (C) concentrations were quantified by multiplex assay. (**D-E**) Relative transcript levels of pro- and anti-inflammatory markers in resident peritoneal macrophages. Each symbol represents an individual mouse. Data are expressed as mean ± SEM. ^*^p<0.05; ^***^p<0.001, unpaired, two-tailed Student’s t-test.

### Hematopoietic SLC37A2 deletion has a minor effect on blood myeloid cell composition in WD-fed Ldlr^-/-^ mice

We next assessed whether hematopoietic SLC37A2 deletion affects monocyte and neutrophil composition in blood as well as in the spleen. **Figures S2A** show our flow cytometry gating strategies. At 9-wk WD feeding, no difference was observed between genotypes regarding the frequency of blood monocytes (CD11b^+^CD115^+^), Gr1^low^ monocytes (CD11b^+^CD115^+^Gr1^low^), Gr1^high^ monocytes (CD11b^+^CD115^+^ Gr1^high^), or neutrophils (CD11b^+^CD115^-^Gr1^+^) (**Figures S2B**), or the ratio of Gr1^low^ and Gr1^high^ monocytes in blood monocytes (**Figures S2C**). After 16 wks of diet feeding, the percentage of blood neutrophils (CD11b^+^CD115^-^Gr1^+^) was significantly increased in the SLC37A2^Δhema^ mice (**Figures S2D**). Despite the elevated plasma cholesterol and increased blood neutrophils, no difference was detected between genotypes in blood monocyte composition at 16-wk diet feeding (**Figures S2D** and **S2E**). We did not observe any significant changes in the monocyte and neutrophil composition in the spleen between genotypes (**Figures S2F** and **S2G**). Overall, these results suggest hematopoietic SLC37A2 deletion has a minimal effect on blood myeloid cell composition. Note that the concentration of plasma MCP-1, a primary chemokine recruiting monocytes and macrophages from bone marrow to circulation or from the blood circulation to tissues, was significantly lower in the SLC37A2^Δhema^ mice (Figure 3B). We reason that the unaltered blood and spleen monocytes may be the net effect of the combination of decreased plasma MCP-1 and increased plasma cholesterol in the SLC37A2^Δhema^ mice. Taken together, our results suggest that hematopoietic SLC37A2 deletion has a minor effect on blood myeloid composition except for a slight increase in blood neutrophils after 16-wk WD feeding.

### Hematopoietic SLC37A2 deficiency promotes atherosclerosis in WD-fed Ldlr^-/-^ mice

Abnormal lipid metabolism and enhanced local and systemic inflammation accelerate atherosclerosis. Since we observed increased plasma lipids, especially apoB containing lipoproteins, and decreased IL-10 in plasma, we next investigated the development of atherosclerosis in SLC37A2^Δhema^ mice compared to controls. We found that hematopoietic SLC37A2-deficient mice showed a 51% increase in aortic root lesions stained with Oil Red O (**Figures 4A-4B**), despite similar CD68 (macrophage marker) (**Figures 4C-4D**), suggesting that hematopoietic SLC37A2 deletion accelerates atherosclerotic plaque formation but has no effect on macrophage content. Additionally, hematopoietic SLC37A2 deletion also did not affect T cell content in the plaque, as shown by similar CD3 (T cell marker) staining between genotypes (**Figures S3**). We then assessed whether there is an association between plasma lipids vs. atherosclerosis or between plasma IL-10 vs. atherosclerosis in diet-fed mice by performing linear regression analysis. Despite no significant association between intimal area and plasma TC (**Figure 4E**), there was a significant inverse correlation between plasma IL-10 and intimal area (**Figure 4F**), suggesting that the attenuated anti-inflammatory response in the SLC37A2^Δhema^ mice may be the primary driver of enhanced atherosclerosis in those mice.

**Figure 4.**
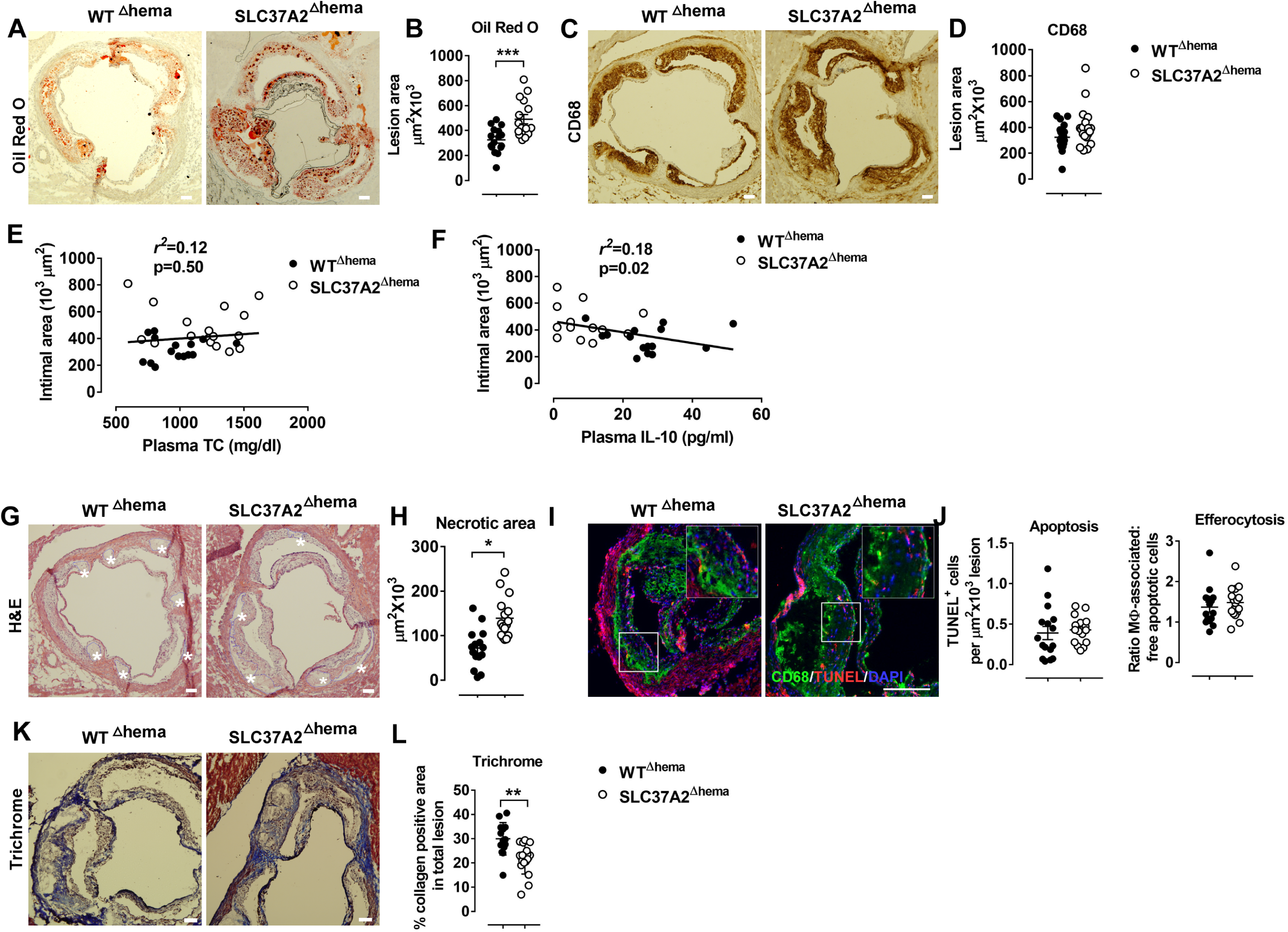
Hematopoietic SLC37A2 deficiency promotes atherosclerosis in WD-fed Ldlr^-/-^ mice. Hematopoietic SLC37A2 knockout (SLC37A2^Δhema^) and control (WT^Δhema^) Ldlr^-/-^ mice were fed a high-fat western diet for 16 wks (n=12-16 mice per genotype). (**A-B**) Quantification of Oil Red O positive intimal area in the aortic root. (**C-D**) Quantification of CD68^+^ cells (macrophages) in the aortic root intimal area. (**E**) Linear regression analysis of aortic root intimal area vs. plasma total cholesterol of 16-wk diet-fed mice. Each point represents an individual animal of the denoted diet group and the correlation coefficient is shown for the entire data set. (**F**) Linear regression analysis of aortic root intimal area vs. plasma IL-10 of 16-wk diet-fed mice. Each point represents an individual animal of the denoted diet group and the correlation coefficient is shown for the entire data set. (**G-H**) Quantification of the aortic necrotic intimal area (total necrotic area) in H&E stained aortic root sections. White stars indicate the necrotic area in the plaques (G). (**I-J**) Quantification of apoptosis (TUNEL-positive cells) and efferocytosis (the ratio of macrophage-associated: free apoptotic cells) in aortic root sections. CD68 (green; macrophages), TUNEL (red), and DAPI (blue; nuclei) were labeled fluorescently. (**K-L**) Quantification of aortic Trichrome stain positive cells. Scale bars = 100 μm. Each symbol represents an individual mouse. Data are expressed as mean ± SEM. n = 12-16 mice per group. ^*^p < 0.05; ^**^p < 0.01; ^***^p<0.001.

IL-10 producing macrophages is responsible for the engulfment and clearance of apoptotic cells (i.e., efferocytosis) ^35^. Failure of efferocytosis leads to pro-inflammatory and immunogenic consequences due to secondary necrosis, exaggerating atherosclerosis ^36,37^. We next examined necrotic core formation and quantified apoptosis and efferocytosis in the plaques. Compared to WT control, SLC37A2^Δhema^ mice had a significant increase (∼40%) in the necrotic area in aortic lesions (**Figure 4G and 4H**). However, no difference was observed regarding the frequency of apoptosis or efferocytosis in lesions between genotypes at 16 wks of diet feeding (**Figure 4I** and **4J**). Interestingly, when we stained the aortic root sections with Masson’s Trichrome stain, we observed a 30% reduction in Trichrome positive staining in SLC37A2^Δhema^ vs. WT control aortic root sections (**Figure 4K** and **4L**), suggesting that hematopoietic SLC37A2 deficiency decreases collagen deposition, likely resulting from impaired alternative macrophage activation. As collagen formation is associates with the stability of plaques ^38,39^, our data suggest that SLC37A2 deficiency in bone marrow promotes plaque instability. Together, our results indicate that hematopoietic SLC37A2 deletion worsens atherosclerosis, which is inversely associated with plasma IL-10 levels.

### Hematopoietic SLC37A2 deletion has minimal impact on insulin resistance and adipose inflammation under pro-atherogenic conditions

In addition to atherosclerosis assessment, we also tested whether hematopoietic SLC37A2 deletion influences adipose tissue inflammation and/or obesity and insulin resistance under pro-atherogenic conditions. WT and SLC37A2^Δhema^ mice gained similar body weight over the 16 wks of diet feeding (**Figures S4A**). Tissue (including liver, spleen, and epididymal fat) mass was comparable between genotypes (**Figures S4B**). Both genotypic mice showed similar glucose clearance and insulin tolerance around 10 wks of diet feeding (**Figures S4C** and **S4D**). As expected, Slc37a2 mRNA expression was significantly reduced in the SLC37A2^Δhema^ vs. control mouse epididymal fat. However, hematopoietic deletion of SLC37A2 did not alter adipose tissue inflammation, as quantified by qPCR analysis of gene expression of macrophage pro- and anti-inflammatory markers, except for increasing ccr2 (MCP-1 receptor) expression (**Figures S4E**). These results suggest that deletion of SLC37A2 in bone marrow cells has minimal impact on WD-induced obesity, insulin resistance, and adipose tissue inflammation under pro-atherosclerotic conditions.

### SLC37A2 deficiency promotes oxLDL-induced macrophage inflammation

OxLDL promotes foam cell formation and triggers oxidative stress and pro-inflammatory responses in macrophages ^40,41^, hence contributing to atherosclerotic plaque formation. Given that there was no significant increase in pro-inflammatory responses in SLC37A2^Δhema^ vs. control mice after 16-wk WD feeding, we asked whether SLC37A2 deletion can affect oxLDL-induced macrophage inflammation *in vitro*. We found that oxLDL markedly increased SLC37A2 protein expression in BMDMs after 24 h stimulation (**Figure 5A**). OxLDL promoted cholesterol, particularly CE, accumulation in macrophages in a dose-dependent manner (**Figure 5B**), regardless of genotypes. However, SLC37A2 deletion does not affect cholesterol loading in macrophages (**Figure 5B**), suggesting a dispensible role of SLC37A2 in macrophage foam cell formation. Like LPS-stimulated cells, *Slc37a2*^*-/-*^ macrophages showed increased expression of Il-1β and Il-6, but not Tnf at the transcriptional level in response to 24 h of oxLDL stimulation (**Figure 5C**), suggesting that SLC37A2 is a stress-responsive protein and increased SLC37A2 expression likely serves as a protective mechanism for resolution of stress-induced inflammation.

**Figure 5.**
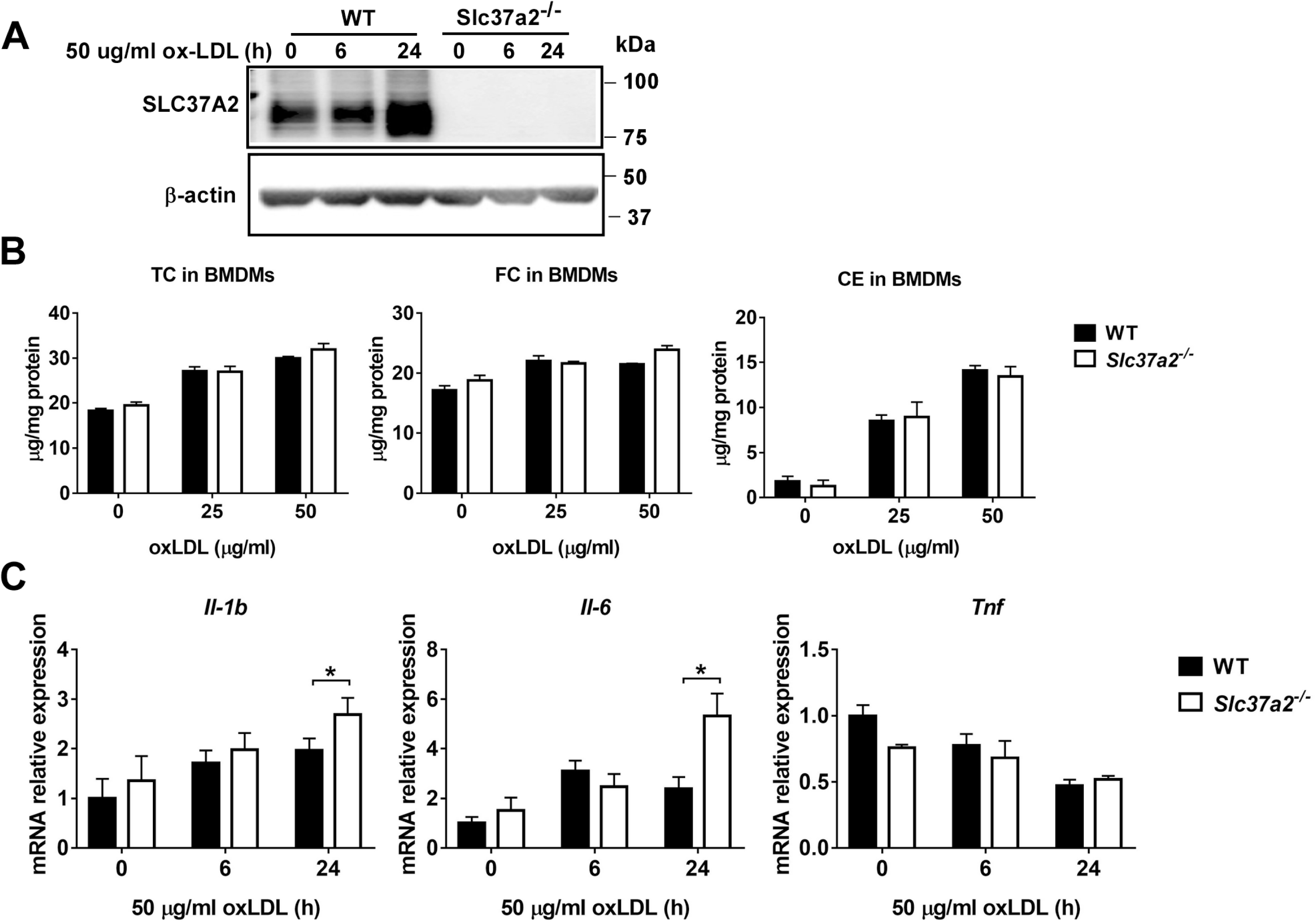
SLC37A2 deficiency promotes oxidized LDL (OxLDL)-induced macrophage inflammation *in vitro*. (**A**) SLC37A2 protein expression analyzed by Western blotting in WT and *Slc37a2*^*-/-*^ bone marrow-derived macrophages (BMDMs) treated with 50 μg/ml oxLDL for 0-24 h. (**B**) Quantification of cellar total cholesterol (TC), free cholesterol (FC), and cholesterol ester (CE) in WT and *Slc37a2*^*-/-*^ BMDMs after treated with or without 25 μg/ml or 50 μg/ml oxLDL for 24 h. (**C**) Relative transcript level of cytokines in WT and *Slc37a2*^*-/-*^ BMDMs stimulated with 50 μg/ml oxLDL for 0-24 h, measured by qPCR. Data are representative of two independent experiments with 3 samples per group (mean ± SEM). ^*^p < 0.05; unpaired, two-tailed Student’s t-test.

### SLC37A2 deficiency impairs M (IL-4) macrophage activation

IL-4 signaling activates STAT6, inducing the expression of genes involved in fatty acid metabolism and transcriptional regulation (such as transcriptional factors PPARs and PPARγ coactivator PGC1β) for reprogramming macrophage lipid metabolism ^42^. We found that IL-4 induced a 1.5 fold increase of Slc37a2 transcript at 6 h (**Figure 6A)** and a 2-fold increase of SLC37A2 protein at 24 h of stimulation (**Figure 6B**) in WT macrophages. Consistent with our *in vivo* findings, SLC37A2-deficient macrophages showed decreased expression of M2 markers, including Arg1, Mrc1, and Ym1 (**Figure 6C**), suggesting that SLC37A2 expression is necessary for M (IL-4) polarization. Interestingly, SLC37A2 deletion also lowered MCP-1 expression in macrophages, consistent with the lower *in vivo* MCP-1 expression in diet-fed SLC37A2^Δhema^ vs. control mice. However, IL-4 does not induce IL-10 expression in either genotypic macrophages, suggesting that M (IL-4) is not a major source of IL-10 in macrophages. Note that WT and SLC37A2 deficient macrophages displayed similar expression levels of metabolic regulators (**Figure 6D**) and fatty acid oxidation genes (**Figure 6E**), which are primarily regulated by STAT6 signaling, suggesting that SLC37A2 deficiency does not impair the STAT6 signaling. Together, our data suggest that SLC37A2 deficiency impairs M (IL-4) polarization, independent of PPARs and PGC1 expression.

**Figure 6.**
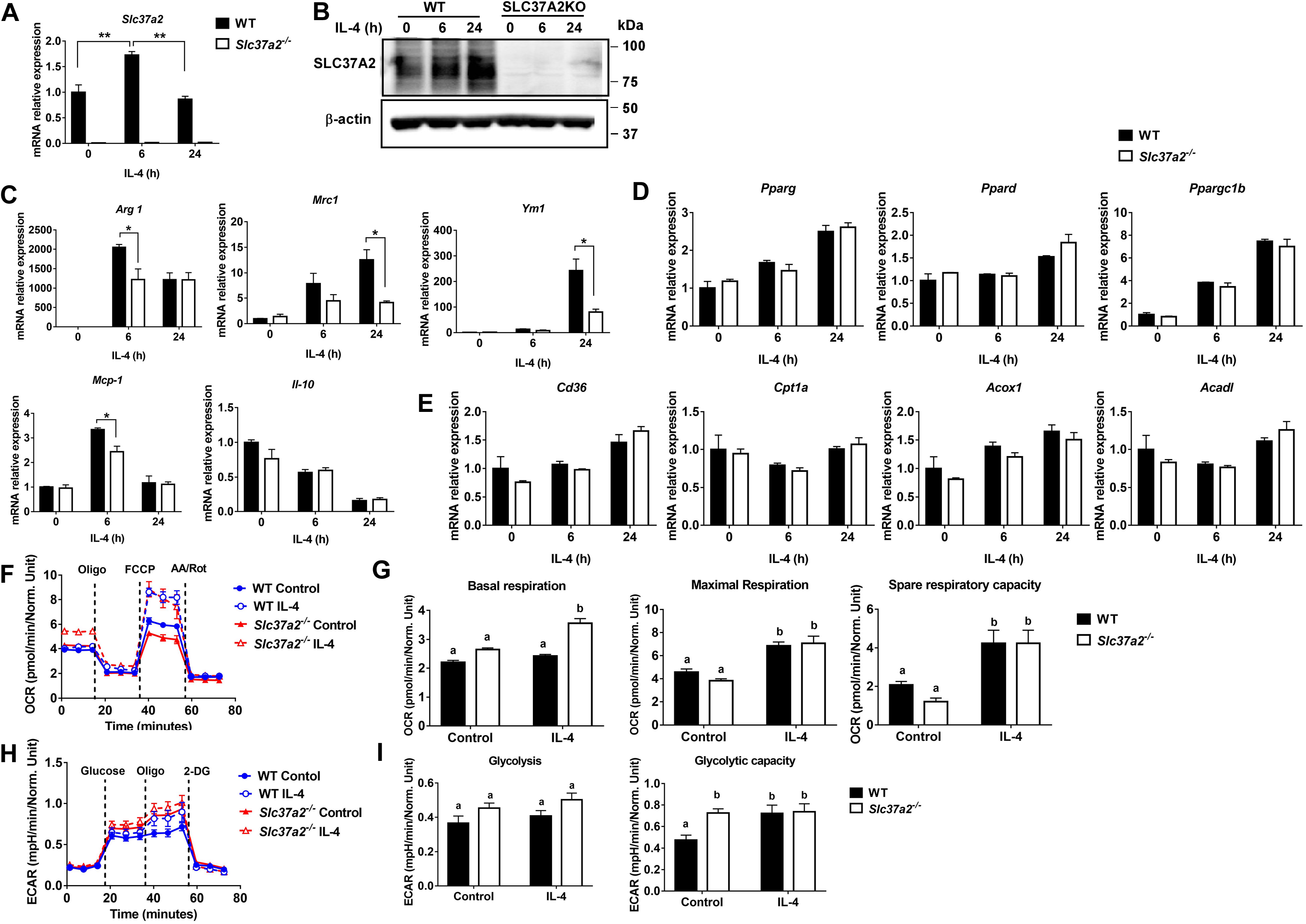
SLC37A2 deficiency impairs IL-4-induced macrophage activation *in vitro*. (**A-B**) SLC37A2 expression in WT and *Slc37a2*^*-/-*^ bone marrow-derived macrophages (BMDMs) stimulated with 20 ng/ml IL-4 for 0-24 h. (A) relative transcript expression and (B) protein expression of SLC37A2 measured by qPCR and western blotting, respectively. (**C**) Relative transcript level of macrophage alternative activation markers, including arginase -1 (Arg), mannose receptor, type I (Mrc1), Ym1, IL-10, and chemokine MCP-1, in WT and *Slc37a2*^*-/-*^ BMDMs stimulated with 20 ng/ml IL-4 for 0-24 h. (**D-E**) The relative transcript level of PPARs and genes encoding transporters or enzymes involved in fatty acid uptake or β-oxidation in WT and *Slc37a2*^*-/-*^ BMDMs stimulated with 20 ng/ml IL-4 for 0-24 h. (**F**) Seahorse analysis of oxygen consumption rate (OCR) in WT and *Slc37a2*^*-/-*^ BMDMs treated with or without 20 ng/ml IL-4 for 24 h. (**G**) Seahorse analysis of extracellular acidification rates (ECAR) in WT and *Slc37a2*^*-/-*^ BMDMs treated with or without 20 ng/ml IL-4 for 24 h. Data are representative of two independent experiments with 3 samples per group (mean ± SEM). ^*^p < 0.05; ^**^p < 0.01; unpaired, two-tailed Student’s t-test (A, C, D, and E). Values with different superscripts differ significantly (p<0.05); two-way ANOVA with post hoc Tukey’s multiple comparisons test (G and I).

Given that mitochondrial oxidative phosphorylation (OXPHOS) supports M (IL-4) macrophage activation ^43^, we next examined mitochondrial respiration by measuring OCR in BMDMs treated with or without 20 ng/ml IL-4 for 6 or 24 h. As expected, 24 h IL-4 stimulation promoted mitochondrial OXPHOS in WT macrophages, as shown by increased maximal OCR and spare respiratory capacity (**Figure 6F**). Interestingly, *Slc37a2*^*-/-*^ vs. WT BMDMs displayed significantly higher basal respiration after 24 h of IL-4 stimulation (**Figure 6F** and **6G**). However, no genotypic difference was observed in maximal respiration or spare respiratory capacity in IL-4 treated cells, suggesting a minor impact of SLC37A2 deletion on mitochondrial OXPHOS in M (IL-4) macrophages. Additionally, SLC37A2 deletion has no effect on mitochondrial respiration in 6 h IL-4 treated macrophages (**Figure S5**). Evidence suggests that glycolysis and glucose utilization are required for M (IL-4) macrophage activation ^43^, while another study indicates that glycolysis is dispensible as long as mitochondrial OXPHOS is intact ^44^. Nevertheless, SLC37A2-deficient macrophages showed slightly increased glycolytic capacity at baseline. No difference in glycolysis or glycolytic capacity was observed between genotypes after IL-4 treatment (**Figure 6H** and **6I**). Overall, our results suggest that SLC37A2 deletion does not significantly impact glycolysis or mitochondrial OXPHOS in M (IL-4) macrophage. Thus, the attenuated M (IL-4) activation in SLC37A2-deficient macrophages is likely independent of these two cellular metabolic processes.

### SLC37A2 deficiency impairs M (AC) macrophage activation

One of the striking changes in the diet-fed SLC37A2^Δhema^ vs. control mice is the 70% reduction of plasma IL-10, a major anti-inflammatory cytokine for cellular homeostasis. Macrophages are a major type of phagocytes that can produce a large amount of IL-10 in response to apoptotic cells ^45^. The clearance of ACs and the suppression of inflammation by IL-10 are required to prevent chronic inflammation and reduce atherosclerosis progression. So, next, we examined apoptotic cell-induced macrophage (M (AC)) activation in WT and SLC37A2-deficient macrophages. We first incubated Jurkat T cells with 1 μM staurosporine for 4 h to induce early apoptosis (Annexin V^+^, PI^-^). Under this condition, 60% of Jurkat T cells undergo apoptosis (**Figure S6**). Compared to WT cells, SLC37A2-deficient macrophages showed a marked reduction in IL-10 expression at both transcriptional level (**Figure 7A**) and protein level (**Figure 7B**) in response to ACs. Engulfment of dead cells has been reported to elevate macrophage fatty acids and mitochondrial β-oxidation, which supports NAD^+^ homeostasis and IL-10 production ^46^. To explore the possible mechanisms of the attenuated M (AC) activation in SLC37A2-deficient macrophages, we first compared the efferocytosis index between genotypes. We found SLC37A2 deletion slightly enhanced engulfment of ACs over a 2-h period (**Figure 7C**). Blockade of phagocytosis by using cytochalasin D (**Figure 7D**) or blockade of fatty acid β-oxidation (**Figure 7E**) slightly reduced AC-induced IL-10 production in both genotypes but did not normalize the differential expression of IL-10 between genotypes. Interestingly, blockade of glycolysis using 2-DG led to a significant reduction of IL-10 in engulfed macrophages in both mouse genotypes (**Figure 7F**), suggesting that glycolysis plays an unappreciated role in AC-induced IL-10 production. More interestingly, the blockade of glycolysis normalized the differential IL-10 secretion between genotypes. Next, we examined mitochondrial respiration and glycolysis by measuring OCR and ECAR, respectively. We found that ACs decreased mitochondrial OXPHOS when engulfed by macrophages regardless of genotypes, as shown by decreased maximal respiration without significant changes in basal respiration (**Figure 7G** and **7H**). Furthermore, ACs promoted aerobic glycolysis in WT macrophages, and SLC37A2 deletion impaired AC-induced glycolysis (**Figure 7I** and **7J)**. No genotypic difference in glycolytic capacity was observed in AC-engulfed macrophages. Together, our results suggest that SLC37A2 positively regulates M (AC) activation through modulation of glycolysis and is likely independent of phagocytosis or fatty acid oxidation.

**Figure 7.**
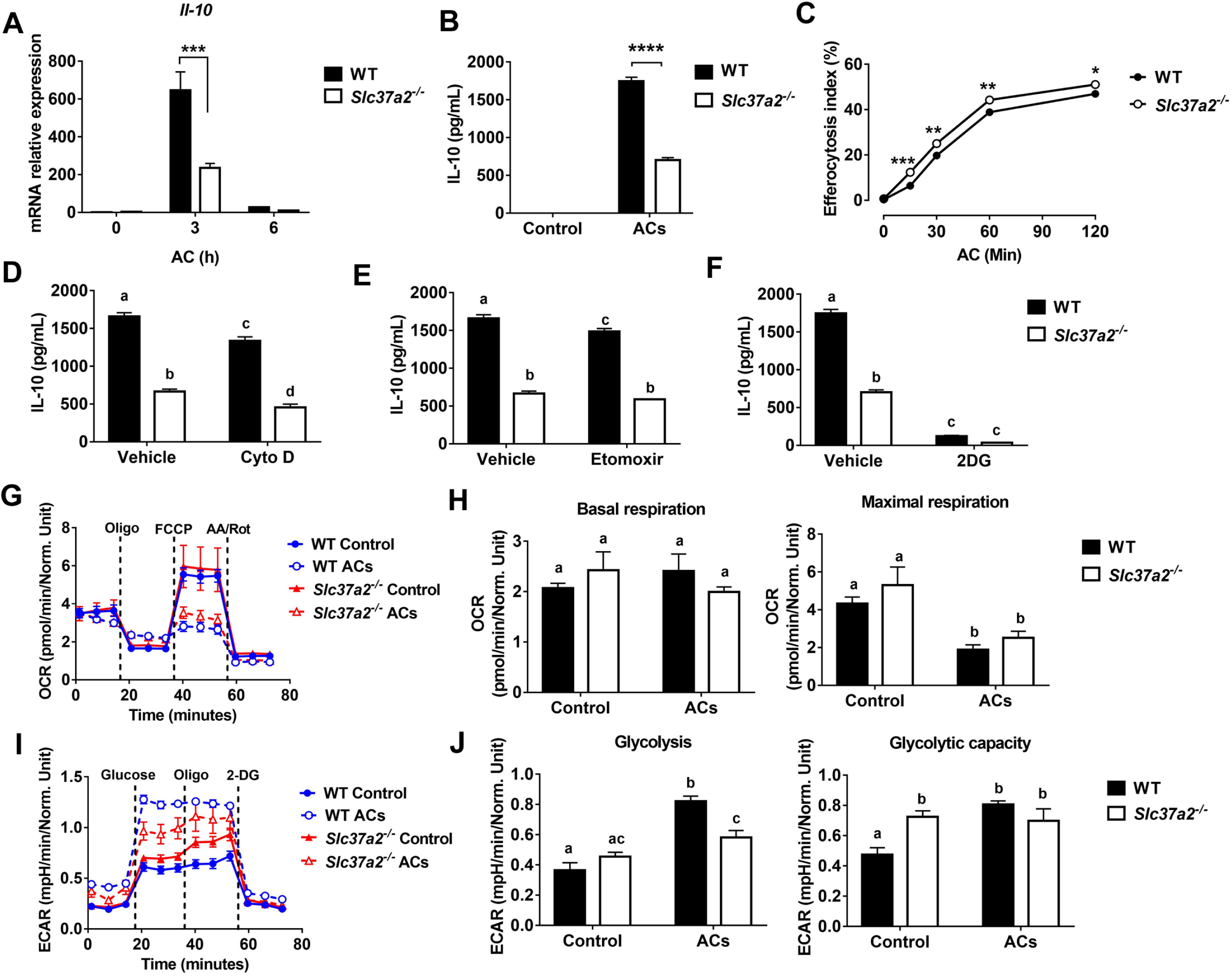
SLC37A2 deficiency impairs apoptotic cell-induced macrophage activation *in vitro*. (**A**) IL-10 transcript expression in WT and *Slc37a2*^*-/-*^ BMDMs stimulated with apoptotic Jurkat T cells (ACs) (macrophages: ACs=1:5) for 0-6 h. Jurkat T cells were incubated with 1 μM staurosporine for 4 h to induce apoptosis. (**B**) IL-10 protein secretion from WT and *Slc37a2*^*-/-*^ BMDMs stimulated with ACs (macrophages: ACs=1:5) for 4 h. (**C**) Efferocytosis index was measured by incubating WT and *Slc37a2*^*-/-*^ BMDMs with ACs (macrophages: ACs =1:5) for 0-120 min. After the indicated time, macrophages were washed with cold PBS 3 times and cell dissociation buffer once before being analyzed by flow cytometry. (**D-F**) IL-10 protein secretion from WT and *Slc37a2*^*-/-*^ BMDMs pretreated with phagocytosis inhibitor cytochalasin D (Cyto D, 10 μM), fatty acid oxidation inhibitor etomoxir (100 μM), and hexokinase inhibitor 2-deoxy-D-glucose (2-DG; 10 mM) for 30 min, followed by co-culture with ACs (macrophages: ACs= 1:5) for 4 h. (**G-H**) Seahorse analysis of oxygen consumption rate (OCR) in WT and *Slc37a2*^*-/-*^ BMDMs treated with or without ACs (macrophages: ACs= 1:5) for 3 h. (**I-J**) Seahorse analysis of extracellular acidification rates (ECAR) in WT and *Slc37a2*^*-/-*^ BMDMs treated with or without ACs (macrophages: ACs= 1:5) for 3 h. Data are representative of two independent experiments with 3 samples per group (mean ± SEM). ^**^ P<0.01, ^***^ P<0.001, ^****^ P<0.0001, unpaired, two-tailed Student’s t-test (A-C). Values with different superscripts differ significantly (p<0.05); two-way ANOVA with post hoc Tukey’s multiple comparisons test (D-J).

## Discussion

As an ER-membrane anchored G6P transporter, SLC37A2 is a critical regulator of LPS-induced macrophage pro-inflammatory activation by modulation of glycolysis ^24^. Herein, we made novel observations that SLC37A2 expression is necessary to maintain IL-4 and apoptotic cell-induced macrophage alternative activation in a glycolysis-independent and –dependently manner, respectively. Hematopoietic expression of SLC37A2 is atheroprotective *in vivo* under pro-atherogenic conditions. Disruption of SLC37A2 significantly impairs macrophage activation induced by IL-4 or apoptotic cells. Moreover, disruption of SLC37A2 in hematopoietic cells impairs anti-inflammatory responses and worsens atherogenesis in high-fat western diet-fed Ldlr^-/-^ mice. Since LPS, oxLDL, and IL-4 induce macrophage SLC37A2 protein expression, we speculate that induction of macrophage SLC37A2 expression promotes inflammation resolution and prevents or slows atherosclerosis progression.

The primary functions of alternatively-activated macrophages are to promote tissue remodeling and repair through collagen formation and clearance of apoptotic cells (efferocytosis). Failure of efferocytosis leads to increased apoptosis and cell death. Alternatively-activated macrophages also secrete high levels of anti-inflammatory cytokines such as IL-10, which is a crucial mediator of inflammation resolution ^47^. IL-10 can also promote efferocytosis by a positive feedback pathway ^48^. Blocking IL-10 accelerates atherosclerosis ^49^, whereas targeting the delivery of IL-10 via nanoparticles attenuates atherosclerosis ^50^. In our study, although no significant changes in apoptosis or efferocytosis in atherosclerotic plaques were observed between genotypes, loss of SLC37A2 in bone marrow did enlarge necrotic cores and decrease collagen deposition in the diet-fed mouse plaques. This could suggest secondary necrosis of apoptotic cells may have nonetheless occurred. Moreover, plasma IL-10, but not plasma cholesterol, shows an inverse correlation with aortic plaque size in diet-fed mice. Because SLC37A2 expression is necessary for anti-inflammatory macrophage activation *in vitro* and *in vivo*, we speculate that the impaired anti-inflammatory responses are the primary driving force of enhanced atherosclerosis in the SLC37A2^Δhema^ mice.

Cellular metabolism has emerged as an essential determinant of macrophage activation in response to microenvironmental cues. Macrophage pro-inflammatory activation was long thought to primarily rely on glucose metabolism, whereas M (IL-4) macrophages switch to fatty acid oxidation and mitochondrial biogenesis to support their anti-inflammatory functions ^42^. However, recent studies challenged this concept and suggested that fatty acid oxidation is dispensable for M (IL-4) macrophage polarization ^51,52^. Whether glycolysis plays a role in M (IL-4) activation is also under debate and requires further investigation ^43^. Therefore, how macrophages reprogram cellular metabolism to favor M (IL-4) polarization remains poorly defined. In our study, SLC37A2 deficiency impairs M (IL-4) polarization independent of PPARs and PGC1. Given that SLC37A2 regulates glycolysis and mitochondrial OXPHOS in LPS-treated macrophages, we hypothesized that SLC37A2 might promote M (IL-4) activation by enhancing mitochondrial respiration. Our results showed that 24 h of IL-4 stimulation significantly increased glycolysis and mitochondrial respiration in macrophages regardless of SLC37A2 expression. Despite the impaired M (IL-4) activation, SLC37A2 deletion slightly increased mitochondrial respiration and had no effect on glycolysis in M (IL-4) macrophages. These results suggest that the impaired M (IL-4) activation in SLC37A2-deficient macrophages is likely regulated by unknown mechanisms rather than rewiring the glycolytic process or altering mitochondrial respiration. Additionally, mitochondrial β-oxidation of fatty acids derived from apoptotic cells has been shown to support efferocytosis-induced IL-10 production ^46^. Interestingly, our study suggests that glycolysis plays a much greater role in AC-induced IL-10 production, as evidenced by the increased ECAR in AC-treated macrophages and a much greater reduction of IL-10 in 2DG-treated macrophages relative to etomoxir-treated cells. Our results agreed with Morioka’s findings that efferocytosis promotes glucose uptake, glycolysis, and lactate production ^53^. Unlike LPS-treated macrophages in which SLC37A2 deletion promotes glycolysis, SLC37A2-deficient macrophages showed attenuated glycolysis in response to AC stimulation. Furthermore, blockade of glycolysis, but not phagocytosis or fatty acid oxidation, normalized the differential secretion of IL-10 between genotypes. Together, our results suggest that glucose metabolism plays a central role in SLC37A2-regulated M (AC) activation.

Our *in vitro* macrophage studies suggest that SLC37A2 deletion enhances pro-inflammatory activation in both LPS- and oxLDL-treated macrophages. However, hematopoietic SLC37A2 deletion does not cause elevated pro-inflammatory responses in WD-fed Ldlr^-/-^ mice. Unexpectedly, plasma MCP-1 showed a 30% reduction in SLC37A2^Δhema^ mice. MCP-1 is one of the critical chemokines that regulate the migration and infiltration of monocytes/macrophages. A higher circulating level of MCP-1 is associated with an increased long-term risk of stroke in the general population ^54^. Inhibition of MCP-1 ^55^ decreases plaque size and limits macrophage infiltration in experimental models of atherosclerosis. As discussed above, hematopoietic SLC37A2 deletion increases lipid deposition and necrotic core formation but does not enhance macrophage (CD68^+^ cells) infiltration in diet-fed Ldlr^-/-^ mice. One possible explanation for the indistinguishable macrophage content between genotypes is the net effect of reducing anti-inflammatory IL-10 and pro-inflammatory MCP-1 in those knockout mice. Interestingly, IL-4 signaling induces MCP-1 expression in macrophages and other cells ^56-58^. We observed a significant decrease in MCP-1 expression induced by IL-4 in SLC37A2-deficient macrophages. Whether the down-regulation of MCP-1 expression results from the impaired IL-4 response in WD-diet fed SLC37A2^Δhema^ mice requires further investigation. On the other hand, macrophage heterogeneity is more complex as activation drives a spectrum of macrophages ^12^. *In vitro* models of macrophage polarization (M1 vs. M2) are more simplified than the microenvironment that macrophages encounter in an atherosclerotic lesion or liver tissue. The mixed pro-inflammatory and anti-inflammatory profile in diet-fed SLC37A2^Δhema^ mice likely reflects a more complex local microenvironment that macrophages encounter in an atherosclerotic lesion or liver tissue in those diet-fed mice.

Alternative activation of hepatic macrophages promotes liver fatty acid oxidation and improves metabolic syndrome ^34^. One interesting observation in the current study is that the SLC37A2^Δhema^ mice displayed increased plasma and liver cholesterol and concomitantly decreased expression of genes encoding liver fatty acid oxidation enzymes and anti-inflammatory macrophage marker expression in the liver after 16-wk WD feeding. These results suggest that SLC37A2 deletion impairs alternative activation of Kupffer cells, leading to decreased fatty acid oxidation and increased neutral lipid accumulation in the liver and plasma. Since Kupffer cell replacement in *Slc37a2*^*-/-*^ BMT mice is incomplete ^59^, the modest elevation of plasma and liver lipids in the SLC37A2^Δhema^ mice may underestimate the harmful effect of SLC37A2 deletion on liver lipid homeostasis in the context of atherosclerosis.

In summary, under *in vivo* pro-atherogenic conditions, hematopoietic SLC37A2 expression is necessary for maintaining alternative macrophage activation and IL-10 production. Loss of hematopoietic SLC37A2 impairs anti-inflammatory activities at the cell (peritoneal macrophages), tissue (liver), and systemic (plasma) levels and accelerates atherosclerosis. Our study suggests that hematopoietic SLC37A2 expression protects against atherosclerosis in mice.

### Limitations of the study

One of the limitations of the BMT model is that BM contains multiple types of immune cells. Many of them, including T cells, B cells, neutrophils, and dendritic cells, are involved in atherogenesis. Because of this, we cannot distinguish the specific contribution of macrophage SLC37A2 from other immune cells to atherosclerosis. Additionally, we only examined atherosclerosis after 16-wks diet feeding, a more advanced stage of atherosclerosis. Lastly, we only used male Ldlr^-/-^ mice as recipient mice to investigate the impact of hematopoietic SLC37A2 deficiency on the pathogenesis of atherogenesis and obesity and insulin resistance induced by high-fat diet feeding. Therefore, we do not know whether there is a sex-dependent effect of hematopoietic SLC37A2 expression on the pathogenesis of atherogenesis or not.

## Nonstandard Abbreviations

AC: apoptotic cells
BMDM: bone marrow-derived macrophages
BMT: bone marrow transplantation
ECAR: extracellular acidification rate
NAD^+^: nicotinamide adenine dinucleotide
OCR: oxygen consumption rate
OXPHOS: oxidative phosphorylation
SLC37A2: solute carrier family 37 member 2
WD: western diet

## Acknowledgments

This study was supported by NIEHS Z01 ES102005 (M.B.F.), NIH R01 HL119962 (J.S.P.), NIH R35 GM126922 (C.E.M.), NIH R01 HL132035 (X.Z.), NIH T32GM127261 (A.K.M.), and NIH T32 AI007401 (A.K.M.). The study was also supported by National Center for Advancing Translational Sciences of the National Institutes of Health under Award Number UL1TR001420 (Research Assistant fund to X.Z.).

