## Supplemental Materials for "Hematopoietic-SLC37A2 deficiency accelerates atherosclerosis in LDL receptor-deficient mice"

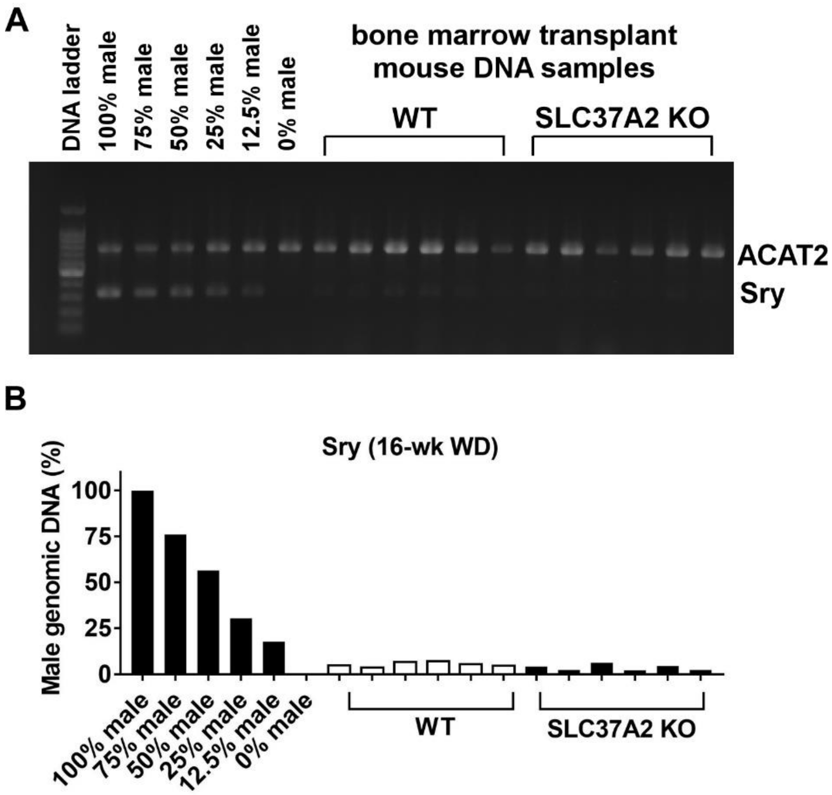


**Figure S1. The efficiency of circulating hematopoietic cell replacement after bone marrow transplantation.**

Sixteen wks after diet feeding, genomic DNA was isolated from whole blood. Fragments of the male Sry gene and ACAT2 (control) genes were amplified by PCR using genomic DNA as a template.

(**A**) A series of male and female genomic DNA mixtures were amplified to construct a standard curve based on the quantified ratio of Sry/ACAT2 band density and the actual percentage of the male/female genomic DNA mixture.

(**B**) Percentage of blood leukocyte female genomic DNA in male recipient mice was calculated based on the standard curve.

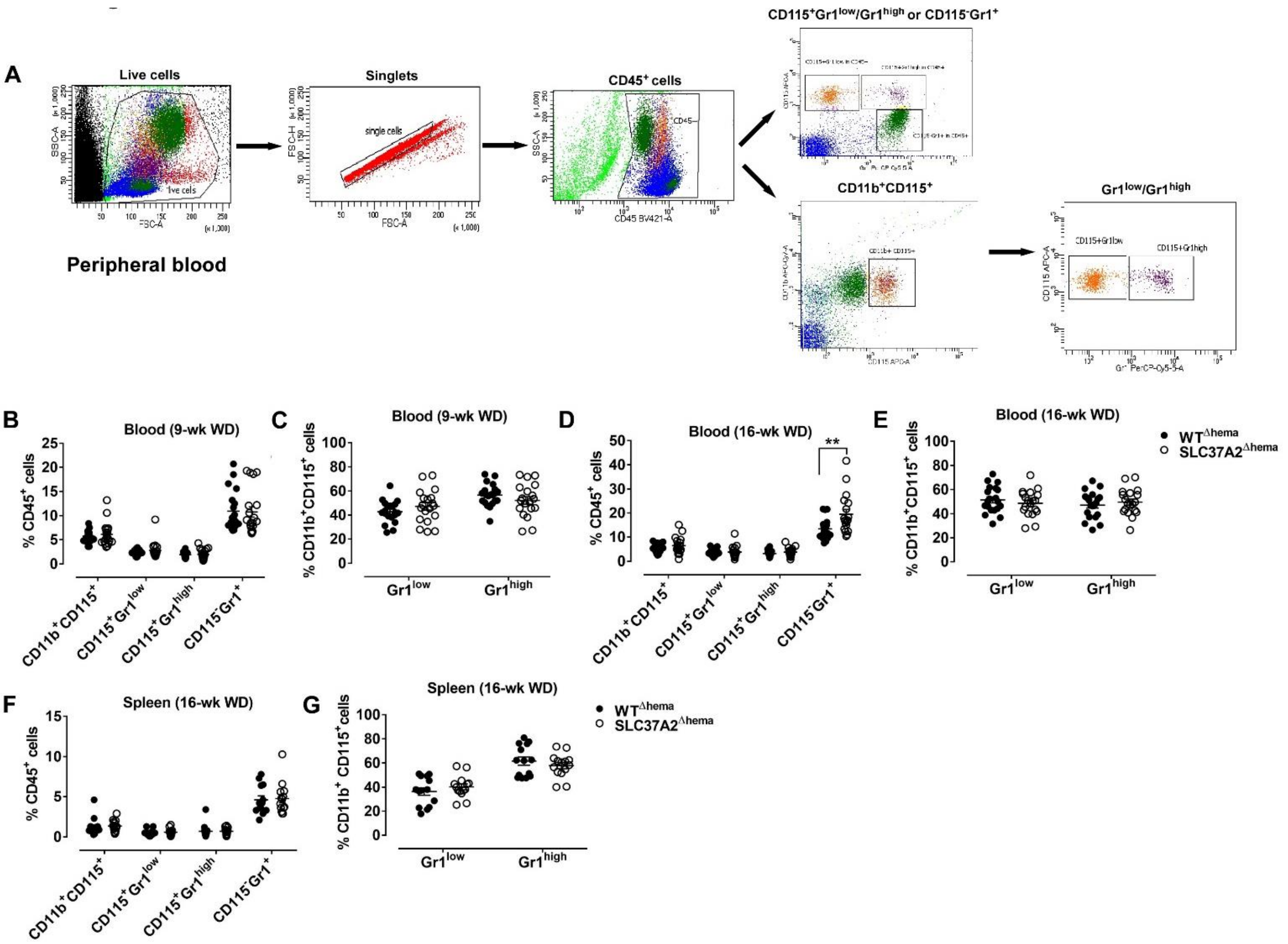


**Figure S2. Hematopoietic SLC37A2 deletion has a minor effect on blood or spleen myeloid cell composition after 16-wks of diet feeding.**

Irradiated Ldlr$^{-/-}$ mice transplanted with WT or SLC37A2KO bone marrow were fed a high-fat western diet for 16 wks. Blood cells and spleen cells were stained with CD11b-PE, CD115-APC, Gr1 (Ly6C/Ly6G)-PerCP-Cy5.5, and CD45-V450 and analyzed by flow cytometry.

(**A**) Gating strategies of the flow cytometry analysis for peripheral blood cells. Similar gating strategies were used to analyze the splenocytes.

(**B-C**) Percentages of monocytes (CD11b$^+$CD115$^+$), Gr1$^{low}$ (CD11b$^+$CD115$^+$Gr1$^{low}$), Gr1$^{high}$ (CD11b$^+$CD115$^+$Gr1$^{high}$), and neutrophils (CD11b$^+$CD115$^-$Ly6G$^+$) in circulating blood CD45$^+$ cell (B) or blood monocytes (C) after 9-wk diet feeding.

(**D-E**) Percentages of monocytes (CD11b$^+$CD115$^+$), Gr1$^{low}$ (CD11b$^+$CD115$^+$Gr1$^{low}$), Gr1$^{high}$ (CD11b$^+$CD115$^+$Gr1$^{high}$), and neutrophils (CD11b$^+$CD115$^-$Ly6G$^+$) in circulating blood CD45$^+$ cell (D) or blood monocytes (E) after 16-wk diet feeding.

(**F**) Percentages of monocytes (CD11b$^+$CD115$^+$), Gr1$^{low}$ (CD11b$^+$CD115$^+$Gr1$^{low}$), Gr1$^{high}$ (CD11b$^+$CD115$^+$Gr1$^{high}$), and neutrophils (CD11b$^+$CD115$^-$Ly6G$^+$) in spleen after 16-wk diet feeding.

Data are expressed as mean ± SEM. n = 12-16 mice per genotype. ** P<0.01, unpaired, two-tailed Student's t-test. Each symbol represents an individual mouse.

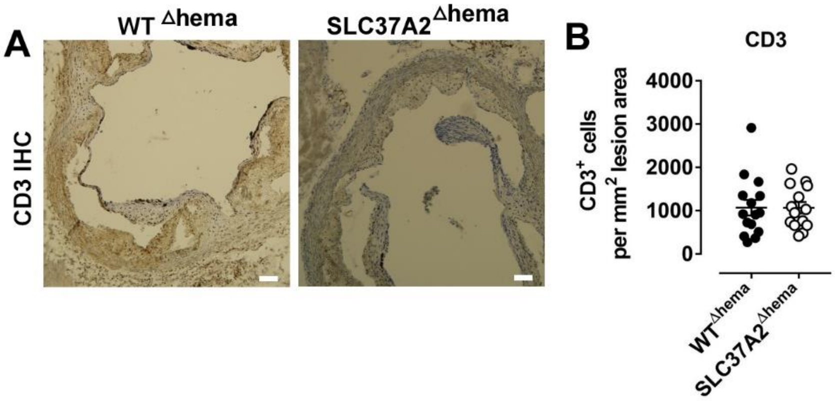


**Figure S3**. **Hematopoietic SLC37A2 deletion does not alter CD3$^+$ cell (T cell) content in the aortic root intimal area.**
Quantification of CD3$^+$ cells (T cells) in the aortic root intimal area (number of CD3 positive cells in lesion). Scale bars = 100 μm.
Data are expressed as mean ± SEM. Each symbol represents an individual mouse. n = 12-16 mice per genotype. Unpaired, two-tailed Student's t-test.

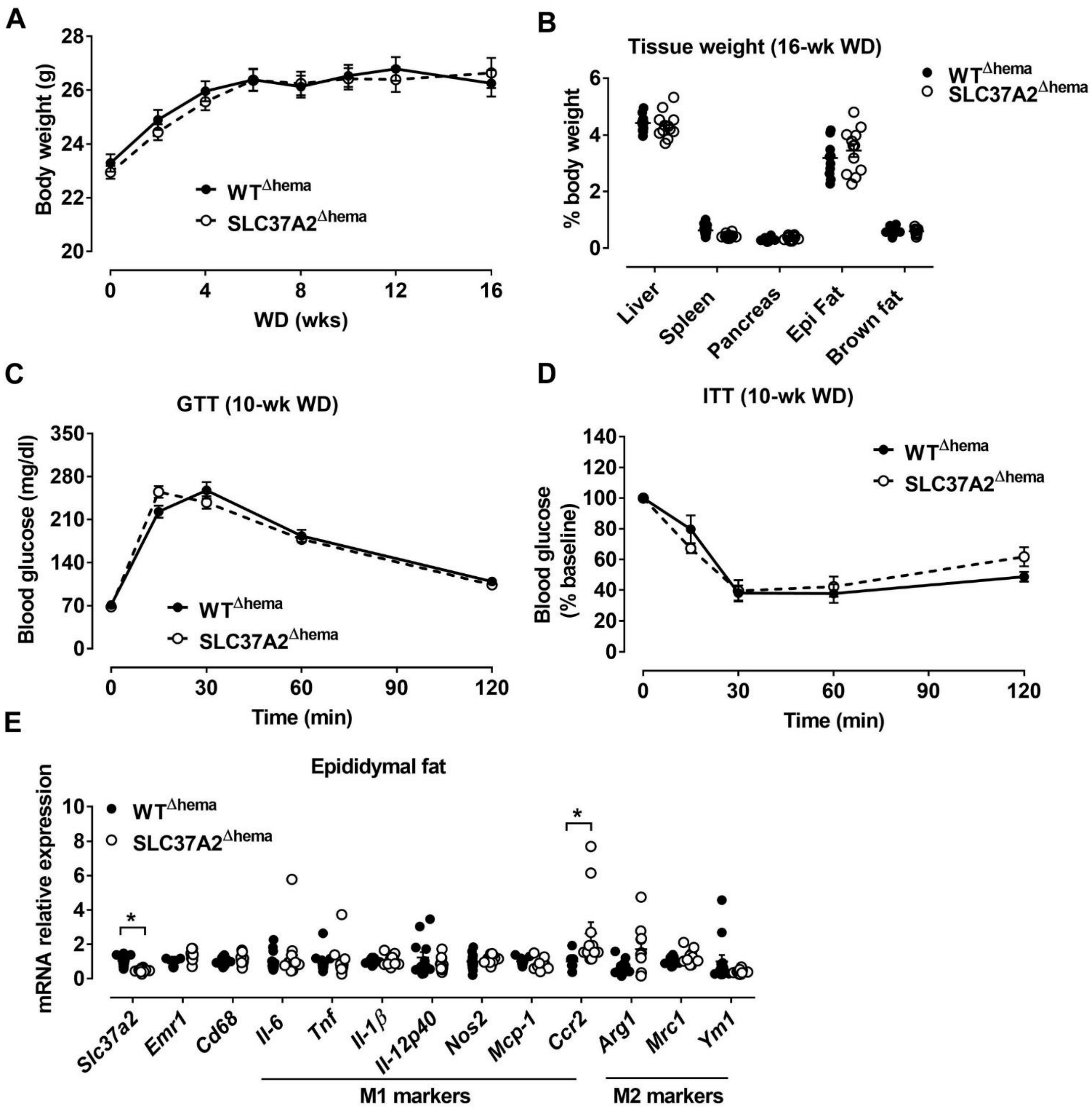


**Figure S4. Hematopoietic SLC37A2 deletion has minimal impact on insulin resistance and adipose inflammation under pro-atherogenic conditions.**
Irradiated Ldlr$^{-/-}$ mice receiving bone marrow from WT or SLC37A2KO mice were fed a high-fat western diet for 16 wks.
(**A**) Mouse body weight over the 16-wk diet feeding period.
(**B**) Mouse tissue weight after 16-wk diet feeding.
(**C**) GTTs were performed after 10-wk diet feeding.
(**D**) ITTs were performed after 11-wk diet feeding.
(**E**) Relative transcript levels of genes in epididymal fat from 16-wk diet-fed mice.

Data are expressed as mean ± SEM. n = 12-16 mice per genotype. * $P<0.05$, unpaired, two-tailed Student's t-test. Each symbol represents an individual mouse.

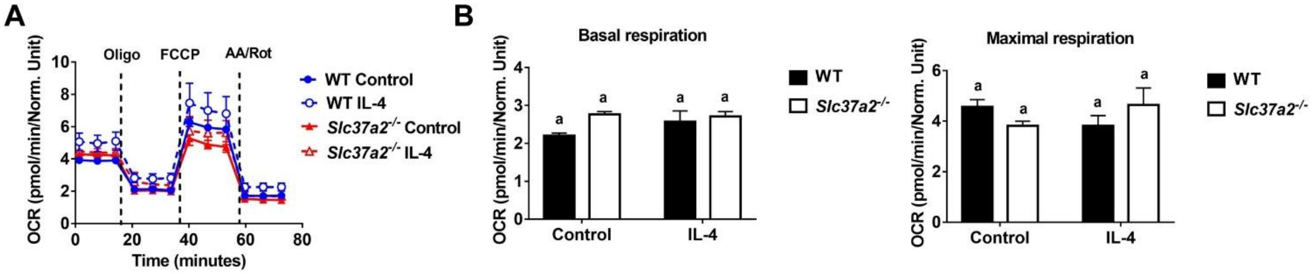


**Figure S5. SLC37A2 deficiency does not affect macrophage mitochondrial respiration after 6 h of IL-4 treatment.**

Seahorse analysis of oxygen consumption rate (OCR) in WT and *Slc37a2*$^{-/-}$ BMDMs treated with or without 20 ng/ml IL-4 for 6 h.

Data are expressed as mean ± SEM. Values with different superscripts differ significantly ($p<0.05$); two-way ANOVA with post hoc Tukey's multiple comparisons test.

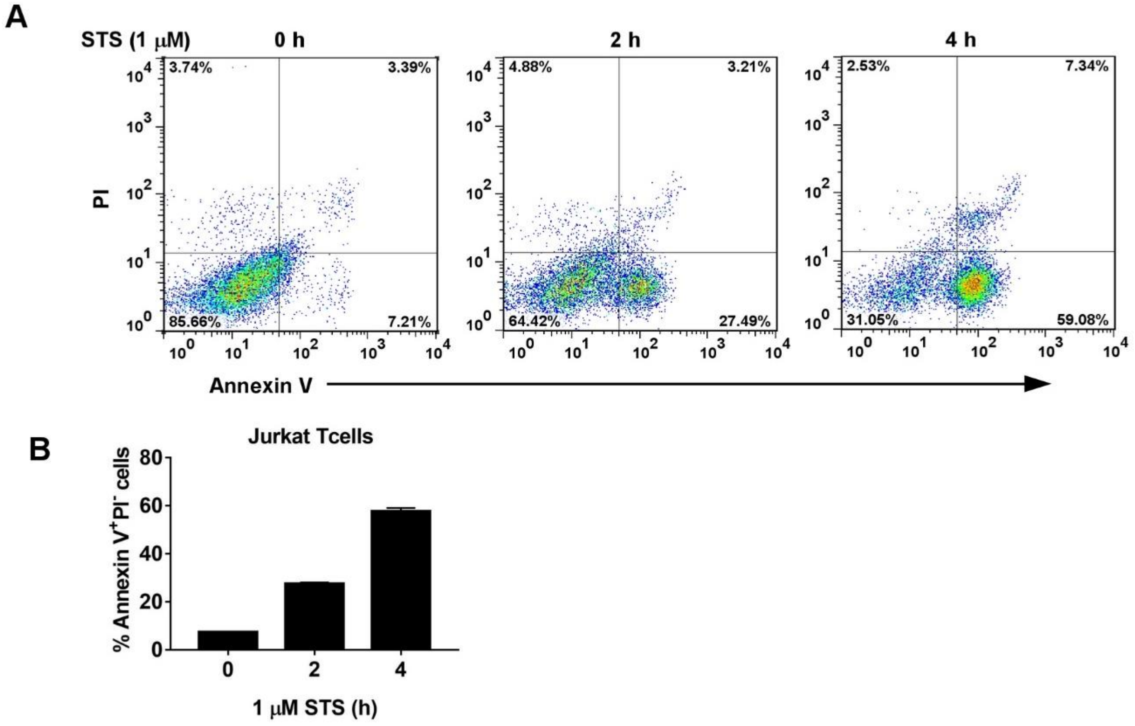


**Figure S6. Flow cytometry analysis of apoptotic Jurkat T cells.**
Jurkat T cells were stimulated with 1 μM staurosporine for 0-4 h before stained with Annexin V-APC and PI.
(**A**) The percentage of early (Annexin $V^{+}PI^{-}$) and late apoptotic (Annexin $V^{+}PI^{+}$) cells was quantified by flow cytometry analysis. The numbers indicate percentages of the subpopulations.
(**B**) The percentage of early apoptotic cells during a 4-h period of staurosporine treatment.

### Table SI. Forward and reverse primers used in qPCR

| Gene name | Forward primer (5'→3') | Reverse primer (5'→3') |
|---|---|---|
| Abca1 | CGTTTCCGGGAAGTGTCCTA | GCTAGAGATGACAAGGAGGATGGA |
| Acox1 | AGATTGGTAGAAATTGCTGCAAAA | ACGCCACTTCCTTGCTCTTC |
| Arg1 | AGCACTGAGGAAAGCTGGTC | CAGACCGTGGGTTCTTCACA |
| Ccr2 | GTGTACATAGCAACAAGCCTCAAAG | CCCCCACATAGGGATCATGA |
| Acadl | TCTTTTCCTCGGAGCATGACA | GACCTCTCTACTCACTTCTCCAG |
| Cd36 | GAGGAATCAGATGAGGATATGGGA | AAGCAGGCTGACTTGGTTGC |
| Cd68 | CTTCCCACAGGCAGCACAG | AATGATGAGAGGCAGCAAGAGG |
| Cpt1a | CACCAACGGGCTCATCTTCTA | CAAAATGACCTAGCCTTCTATCGAA |
| Emr1 | CTTTGGCTATGGGCTTCCAGTC | GCAAGGAGGACAGAGTTTATCGTG |
| Fasn | GCTGCGGAAACTTCAGGAAAT | AGAGACGTGTCACTCCTGGACTT |
| Gapdh | TGTGTCCGTCGTGGATCTGA | CCTGCTTCACCACCTTCTTGAT |
| Hmgcs | GCCGTGAACTGGGTCGAA | GCATATATAGCAATGTCTCCTGCAA |
| Hmgcr | CTTGTGGAATGCCTTGTGATTG | AGCCGAAGCAGCACATGAT |
| Il-1β | GTCACAAGAAACCATGGCACAT | GCCCATCAGAGGCAAGGA |
| Il-6 | CTGCAAGAGACTTCCATCCAGTT | AGGGAAGGCCGTGGTTGT |
| Il-10 | CAGAGCCACATGCTCCTAGA | TGTCCAGCTGGTCCTTTGTT |
| Il-12p40 | AGACCCTGCCCATTGAACTG | GAAGCTGCTGCTGTTCTCATATT |
| Mcp-1 | TTC CTCCACCACCATGCAG | CCAGCCGGCAACTGTGA |
| Mrc1 | ACGAGCAGGTGCAGTTTACA | ACATCCCATAAGCCACCTGC |
| Nos2 | GCAGCTGGGCTGTACAAA | AGCGTTTCGGGATCTGAAT |
| Pparc1a | AACCACACCCACAGGATCAGA | TCTTCGCTTTATTGCTCCATGA |
| Pparc1b | CGCTCCAGGAGACTGAATCCAG | CTTGACTACTGTCTGTGAGGC |
| Ppard | ACGCACCCTTTGTCATCCA | TTCCACACCAGGCCCTTCT |
| Scd1 | CCGGAGACCCCTTAGATCGA | TAGCCTGTAAAAGATTTCTGCAAACC |
| Slc37a2 | GCCTGCGGCAGAAGCAGTGG | AGCAGGGGTGGCCCATGTTG |
| Srebp1c | GGAGCCATGGATTGCACATT | GGCCCGGGAAGTCACTGT |
| Srebp2 | GCGTTCTGGAGACCATGGA | ACAAAGTTGCTCTGAAAACAAATCA |
| Tnf | GGCTGCCCCGACTACGT | ACTTTCTCCTGGTATGAGATAGCAAAT |
| Ym1 | AAGAACACTGAGCTAAAAACTCTCCT | GAGACCATGGCACTGAACG |
